# An evaluation of Nephrology Literature for Transparency and Reproducibility Indicators: Cross-sectional Review

**DOI:** 10.1101/756486

**Authors:** Ian A. Fladie, Tomi Adewumi, Nam Vo, Daniel Tritz, Matt Vassar

## Abstract

**Background:** Reproducibility is critical to diagnostic accuracy and treatment implementation. Concurrent with clinical reproducibility, research reproducibility establishes whether the use of identical study materials and methodologies in replication efforts permit researchers to arrive at similar results and conclusions. In this study, we address this gap by evaluating nephrology literature for common indicators of transparent and reproducible research.

**Methods:** We searched the National Library of Medicine catalog to identify 36 MEDLINE-indexed, English language nephrology journals. We randomly sampled 300 publications published between January 1, 2014, and December 31, 2018. In a duplicated and blinded fashion, two investigators screened and extracted data from the 300 publications.

**Results:** Our search yielded 28,835 publications, of which we randomly sampled 300 publications. Of the 300 publications, 152 (50.67%) were publicly available whereas 143 (47.67%) were restricted through paywall and 5 (1.67%) were inaccessible. Of the remaining 295 publications, 123 were excluded because they lack empirical data necessary for reproducibility. Of the 172 publications with empirical data, 43 (25%) reported data availability statements, 4 (2.33%) analysis scripts, 4 (2.33%) links to a protocol, and 10 (5.81%) were pre-registered.

**Conclusion:** Our study found that reproducible and transparent research practices are infrequently employed by the nephrology research community. Greater efforts should be made by both funders and journals, two entities that have the greatest ability to influence change. In doing so, an open science culture may eventually become the norm rather than the exception.

## Introduction

Reproducibility is critical to diagnostic accuracy and treatment implementation. In nephrology, a substantial body of literature is devoted to establishing the reproducibility of diagnostic tests or procedures. Examples include an evaluation of the reproducibility of the Banff classification for surveillance renal allograft biopsies among pathologists across transplant centers^1^, a novel analytic technique for renal blood oxygenation level-dependent MRI^2^, and a food frequency questionnaire among patients with chronic kidney disease^3^. This form of reproducibility is important clinically, as such studies establish our confidence in tests or procedures for applications to patient care.

Concurrent with clinical reproducibility, research reproducibility establishes whether use of identical study materials and methodologies in replication efforts permit researchers to arrive at similar results and conclusions. In other cases, reproducibility may mean attempts to reanalyze study data to determine whether the same results can be obtained. The National Institute of Diabetes and Digestive and Kidney Diseases (NIDDK) supports the National Institutes of Health’s (NIH) rigor and reproducibility initiative, which was created to foster greater reproducibility of studies funded by taxpayer dollars. The NIDDK also sponsors dkNET^4^, a portal for the dissemination of research protocols and data sets as well as tools and training to promote compliance with the NIH initiative for rigorous and reproducible research^5,6^. Further, the NIDDK directly supports reproducible research through grant funding. As one example, the NIDDK has cosponsored a funding opportunity with other NIH institutes and centers to develop novel, reliable, and cost-effective methods to standardize and increase the utility and reproducibility of human induced pluripotent stem cells. The NIDDK has specifically tasked researchers to develop these stem cells for the replacement of endocrine cells, disease modeling, treatments for diabetic wounds, and reversal of diabetic neuropathy^7^. Research into stem cells has provided significant medical advancements and has the opportunity to demonstrate the importance of reliable and reproducible clinical and basic science research.

While efforts have been made with various stakeholders to foster reproducible research, little is known about the practices actually implemented by researchers involved in nephrology research. In this study, we address this gap by evaluating nephrology literature for common indicators of transparent and reproducible research. By assessing the current state of affairs, we can identify areas of greatest need and establish baseline data for subsequent investigations.

## Methods

### Study design

Our cross-sectional study used a similar methodology as Hardwick *et al.*^*9*^ with our own modifications to evaluate indicators of reproducibility and transparency. Given that this study did not use human subjects, it was not subject to institutional review board approval. When applicable, the Preferred Reporting for Systematic Reviews and Meta-Analyses (PRISMA) guidelines were utilized^10^. The following materials can be accessed on Open Science Framework (https://osf.io/n4yh5/): protocols, raw data, training recording, and additional material.

### Journal and Publication Selection

Nephrology medicine journals were searched on the National Library of Medicine (NLM) catalog on June 5, 2019 by DT for the subject term tag “Nephrology[ST]”. In order to be included in the study, journals had to be MEDLINE indexed, full-text and published in English. To be included in the study, journals also needed to have an electronic International Standard Serial Number (ISSN) and if the electronic ISSN isn’t available, they needed to have a linking ISSN (https://osf.io/tck6m/). DT searched PubMed using the list of ISSN to encompass articles from January 01, 2014 through December 31, 2018. 300 publications were then randomly selected to be included in the analysis. (https://osf.io/mzj45/).

### Data Extraction Training

On June 10, 2019, we had an in person training session led by DT for investigators (TA, IF and NV) on how to extract data. In training, we reviewed study data extraction, design and protocol. As an example, TA, IF and NV extracted data from two publications and reconciled discrepancies after extraction as an example of the process. At the end of the training session, the investigators also applied the same system for the next ten publications to ensure that the process was well standardized and reliable. Starting on July 11, TA, IF and NV conducted extraction of the remaining 289 publications using a duplicate and blinded method. Upon completion of data extraction, the investigators (TA, IF and NV) met to reconcile the discrepancies from data extraction. DT was also available to arbitrate situations when a consensus couldnt be reached among the investigators (TA, IF and NV). The training session from June 10, 2019 was recorded and made available online to investigators for reference (https://osf.io/tf7nw/).

### Data Extraction

We used a Google Form similar to the form used by Hardwicke *et al.* with modifications (https://osf.io/3nfa5/).^9^ Our form contained the following modifications: 5-year impact factor, impact factor for the most recent year listed, additional study design options (cohort studies, case series, secondary analyses, chart reviews, and cross-sectional studies), and additional funding options (university, hospital, public, private/industry, or non-profit). The Google form prompted investigators to assess the overall reporting of transparency and reproducibility characteristics. Data extraction was dependent upon the study design of the publication. Publications without any empirical data were excluded because they fail to provide reproducibility related characteristics. The following study designs were modified: Systematic reviews, meta-analyses, case studies, and case series. Systematic reviews and meta-analyses generally do not contain data measuring materials thus we excluded them from evaluating for material availability. Case reports and case series contain empirical data, but are generally not descriptive enough in their design to be reproduced in subsequent publications and were not expected to contain reproducibility characteristics.^11^

### Open Access Article Availability

We used a systematic approach, to determine if publication were made openly available to the public. First, we searched each publication by title and DOI by using the Open Access Button (https://openaccessbutton.org/). If the Open Access Button failed to identify the publication online or reported an error, we then searched PubMed and Google to identify any other forms of public availability. If the first and second step failed to find the full text, then the publication was determined to be paywall restricted and not available through open access.

### Replication Attempts and Use in Research Synthesis

Using the publication title and the DOI, we searched Web of Science (https://webofknowledge.com) for the following: (1) the number of times a publication was cited by a systematic review/meta-analysis and (2) the number of times a publication was cited by a validity/replication study.

### Statistical analysis

We used Microsoft Excel functions to provide our statistical analysis including percentages, fractions, and confidence intervals.

## Results

We identified 36 nephrology journals that met our inclusion criteria. Our search yielded 28835 publications within our time frame. We randomly sampled 300 publications to include for data extraction. Of the 300 publications, 295 were accessible and contained information to be analyzed. Five publications were inaccessible and therefore were excluded. From the remaining 295 publications, 123 lack empirical data to be analyzed, including 21 that were case studies or case series, therefore they were excluded from our final analysis. Our final analysis included 172 publications with empirical data (Figure 1). Table 1 provided additional information for each indicator used to assess reproducibility and transparency.

**Table 1:**
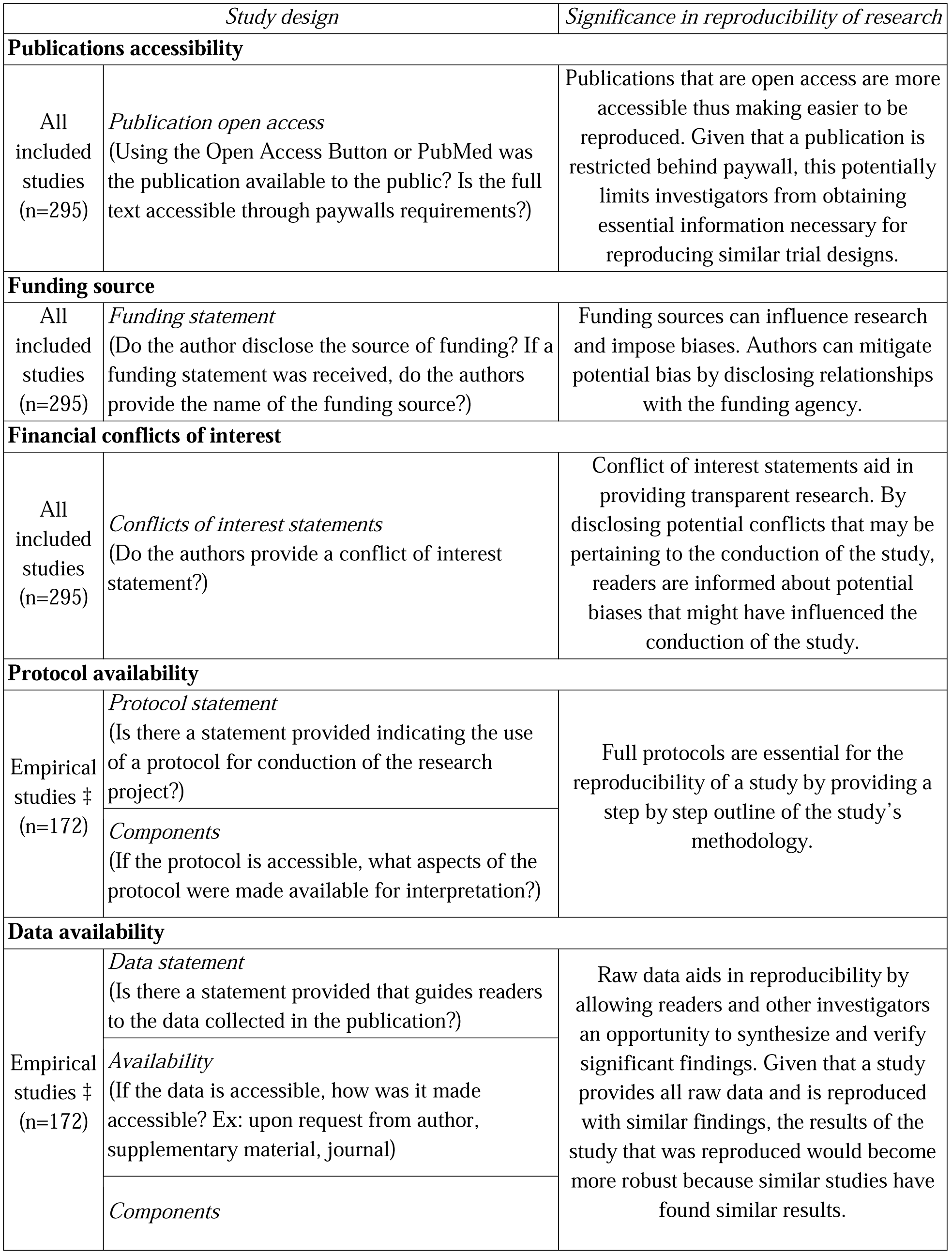
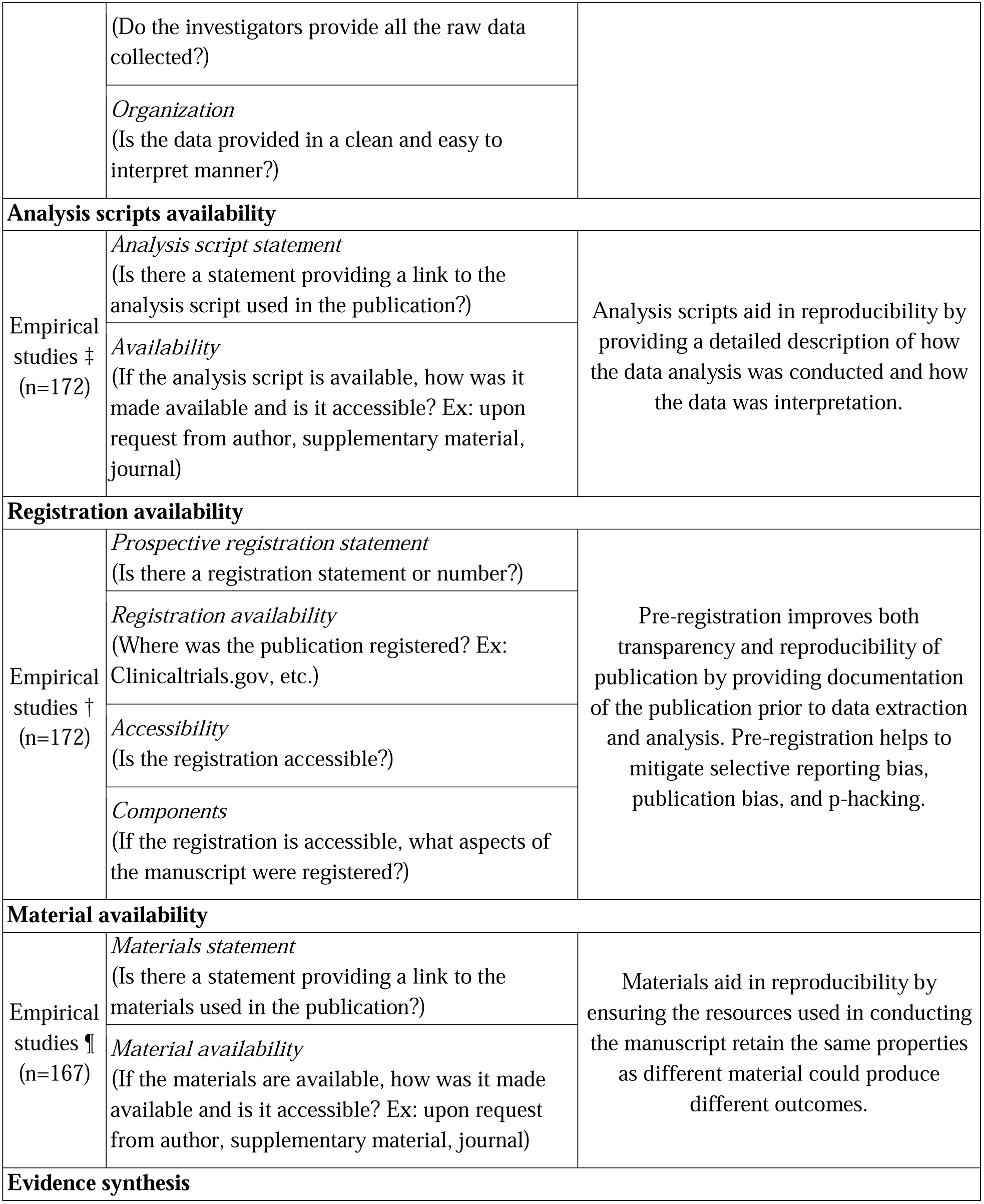
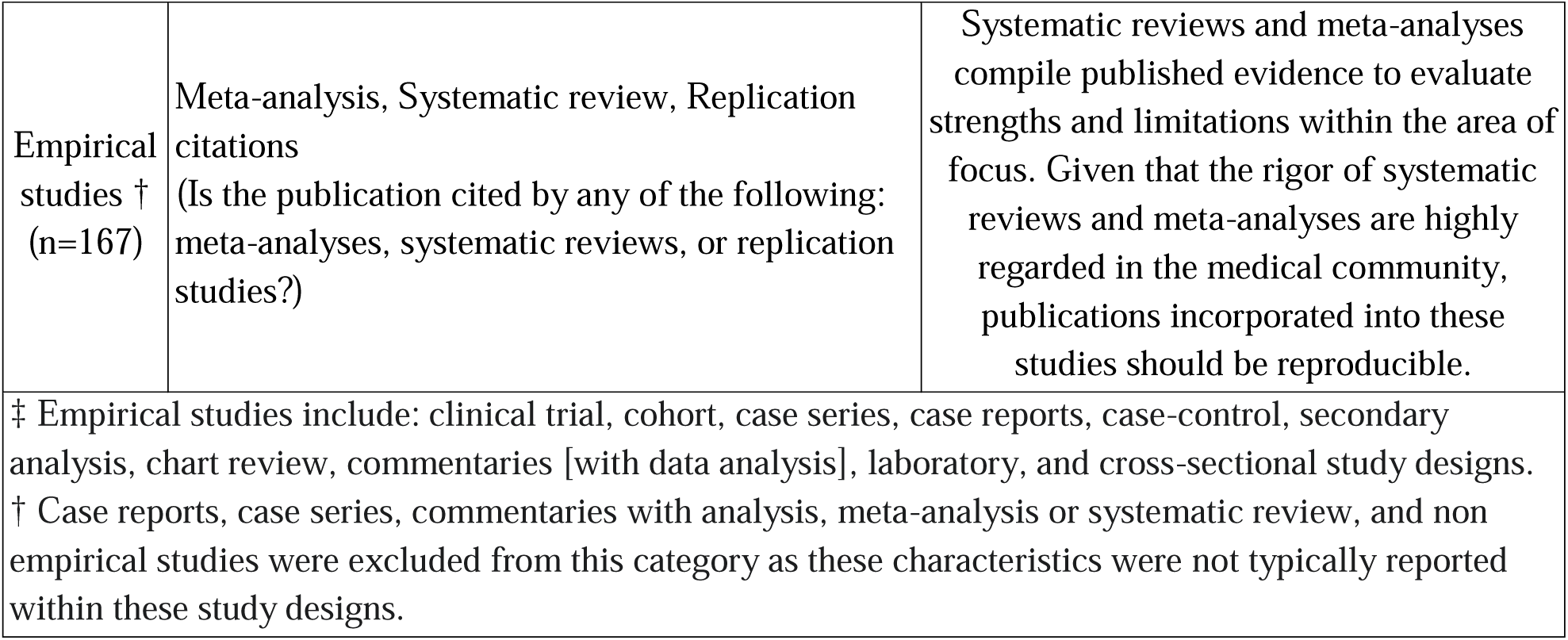
Analyzed components of each publication. Components analyzed varied by study type. Additional details can be found at: https://osf.io/tck6m/

**Figure 1:**
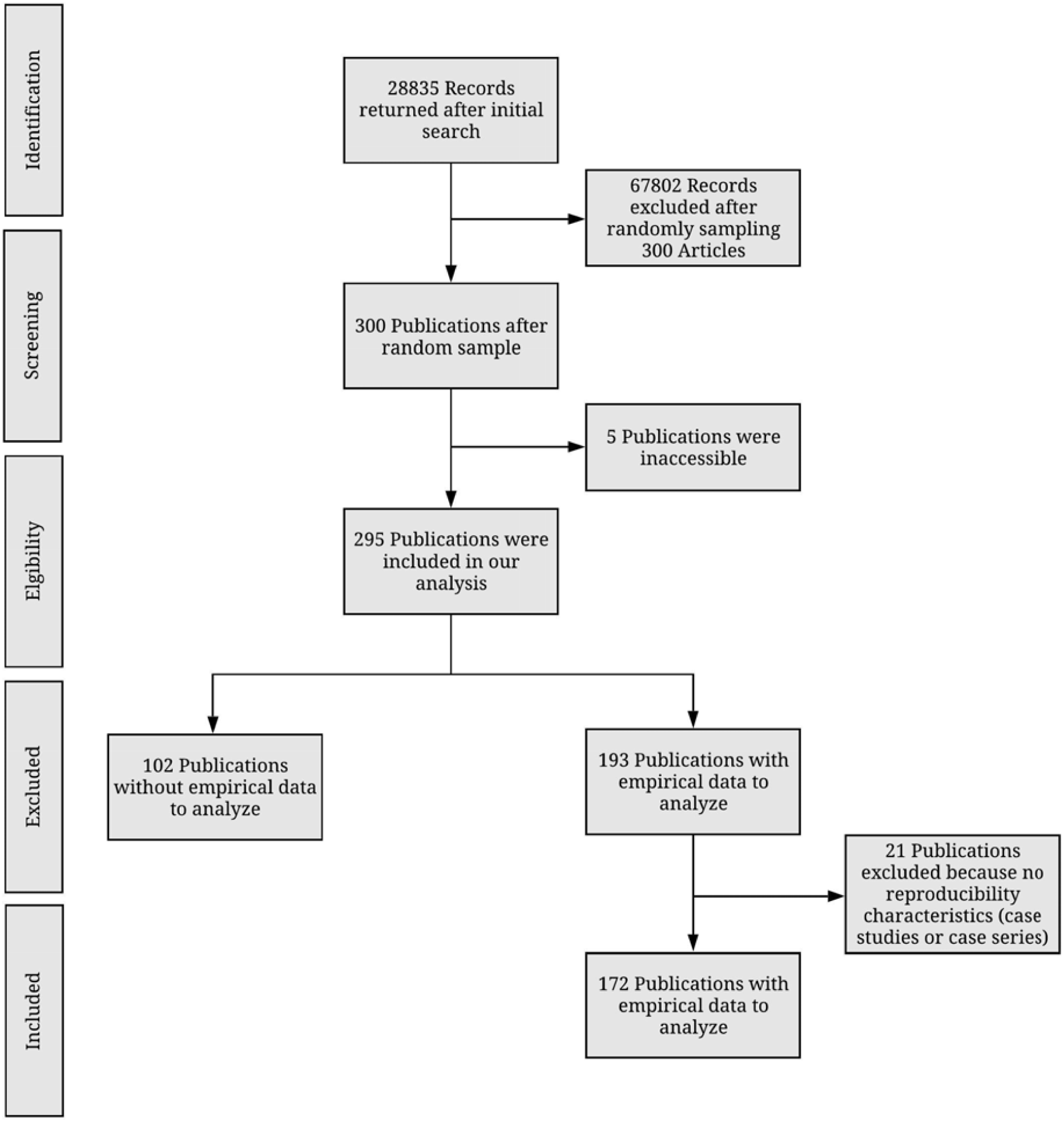
Flow Diagram of included and excluded studies in nephrology journals.

### Sample characteristics

The majority of our publications included in our analysis were cohort (46/295; 15.59%) and Laboratory studies (46/295; 15.59%). Among the 295 publications, the impact factor could not be found for 19 publications. The median 5-year impact factor was 3.232 (IQR 2.053-7.065) with 182 (of 295; 61.69%) of the publications published in United States journals. Furthermore, most corresponding authors were from the United States (97/295; 32.88%).

### Transparency related characteristics

Among the 295 publications that were accessible, each were analyzed transparency characteristics: open access availability, conflict of interest statements, and funding statements. Of these publications, 152 (51.53%) were made open access to the public with the remaining 143 (48.47%) accessible through paywall. Of the 295 publications, 253 (85.76%) provided conflict of interest statements. The majority declared none of the authors had a conflict of interest 188 (63.73%). Approximately one-fifth of publications were funded by public (63/295; 21.34%) whereas hospital (2/295; 0.68%) contributed the least to funding. Furthermore, 19 (of 295; (6.44%) publications reported no funding received to assist in conduction of the publication. Additional transparency characteristics can be found in Table 2.

**Table 2:**
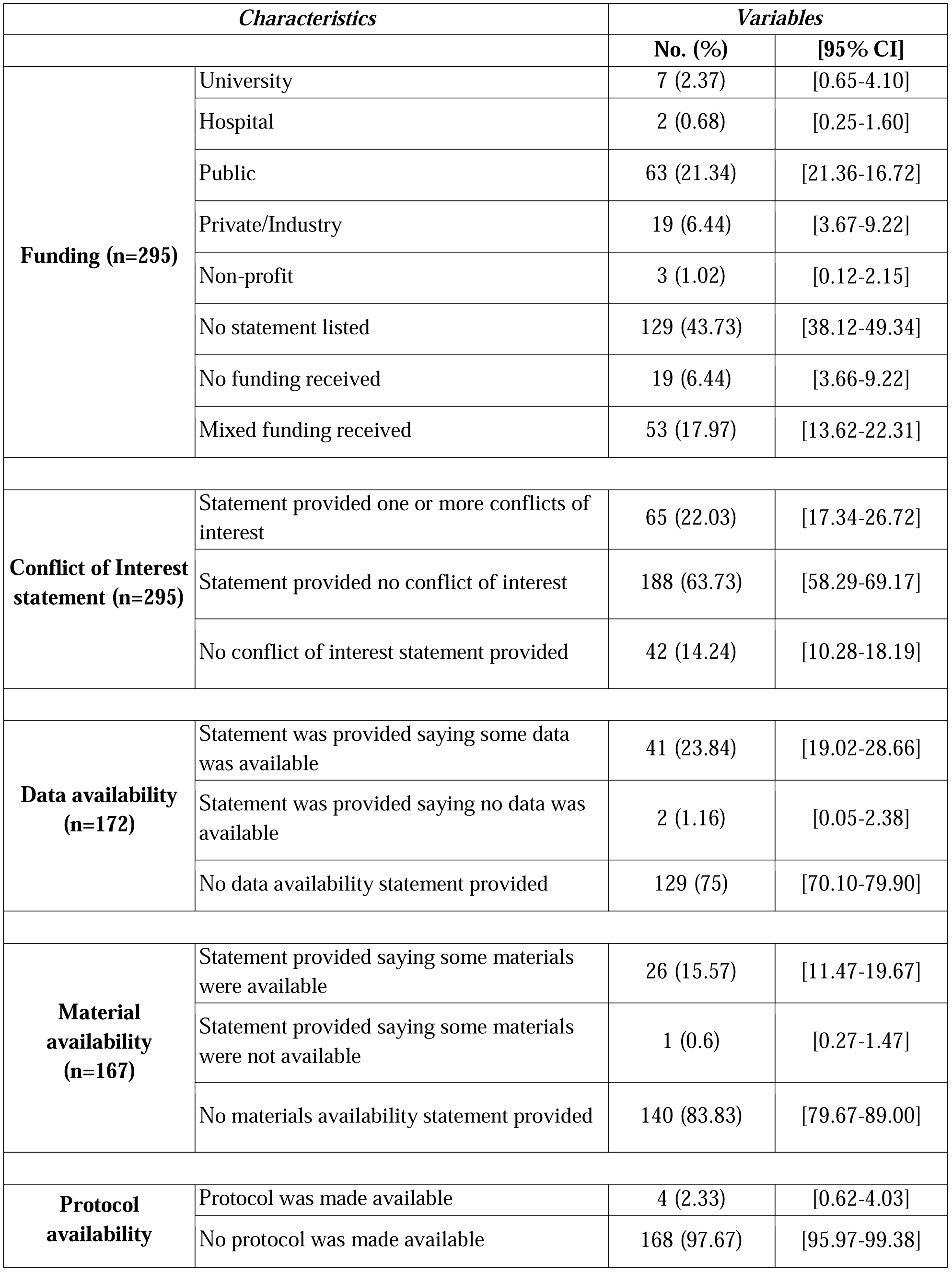
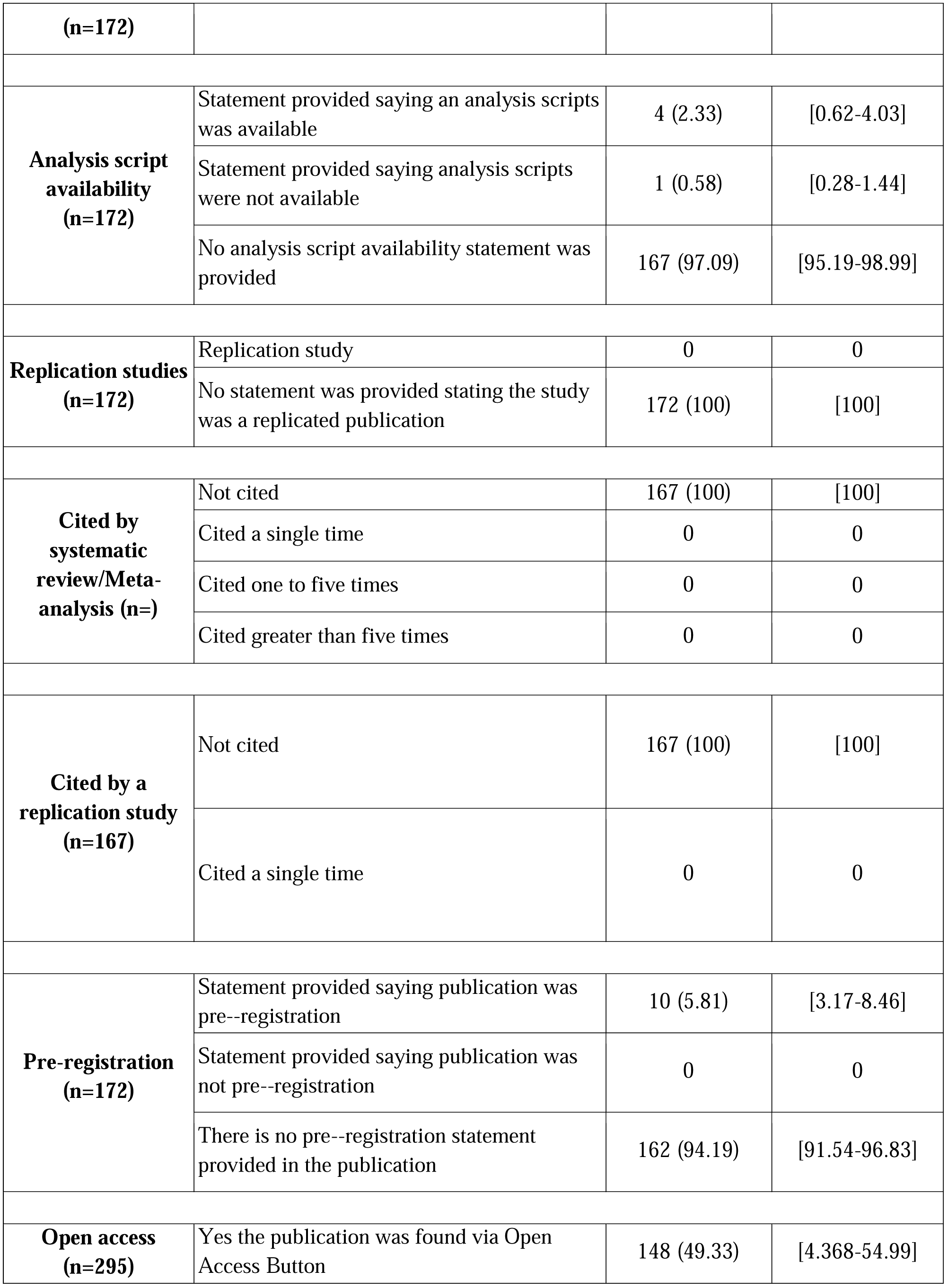
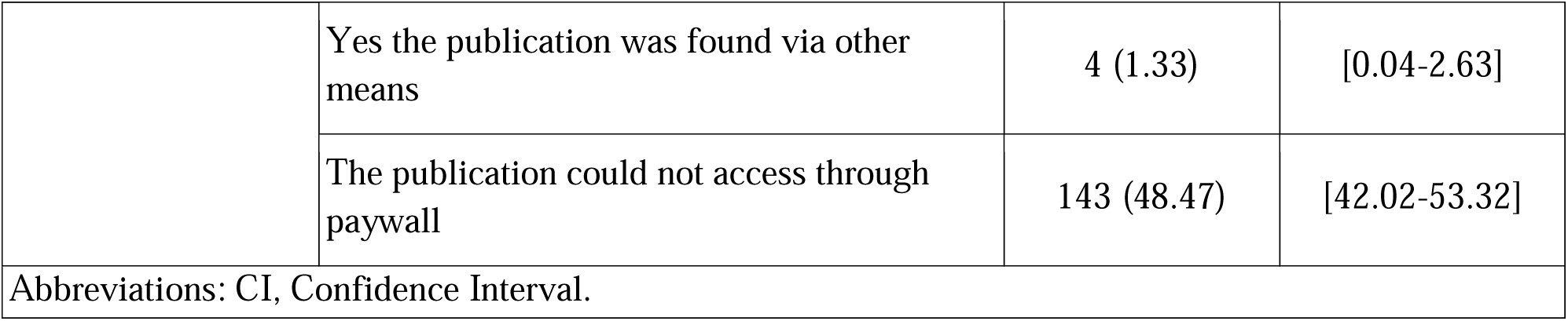
Reproducibility Indicators of Analyzed Orthopaedic Articles.

### Reproducibility related characteristics

A total of 172 publications were analyzed for data, analysis scripts, protocols, preregistration, and material statements. Of these 172 publications, 43 (25%) provided a statement regarding the data used in conducting the trial. Furthermore, few studies were accessible and contained all the raw data used in the publication. The least reported reproducibility characteristic were analysis scripts and protocols (analysis scripts) with only 4 (2.33%) publications containing a statement. Pre-registered studies aid in providing documentation of methods, protocols, analysis scripts and hypotheses prior to data extraction. Among our 172 publications, 10 (5.81%) were pre-registered whereas 162 (94.19%) were not pre-registered. Furthermore, of the publications that were pre-registered, 8 publications contained information regarding the methods. For analysis of materials and evidence synthesis, meta-analysis (n=4), and commentaries with analysis (n=1) were excluded because they lack materials necessary for reproducibility. Of the remaining 167 publications, the majority failed to report material availability statements (140/167; 83.83%). Detailed reproducibility indicator descriptions can be found in Table 2 and Table 3.

**Table 3:** Additional Reproducibility Characteristics.

| **Characteristics** |  | **Variables** |
| --- | --- | --- |
|  |  | **No. (%)** |
| **Type of study**<br>**(n=295)** | No empirical | 102 (34.58) |
|  | Meta-analysis | 4 (1.36) |
|  | Commentary with analysis | 1 (0.34) |
|  | Clinical trial | 17 (5.76) |
|  | Case study | 18 (6.10) |
|  | Case series | 3 (1.02) |
|  | Cohort | 46 (15.59) |
|  | Chart review | 16 (5.42) |
|  | Case control | 4 (1.36) |
|  | Data survey | 8 (2.71) |
|  | Laboratory | 46 (15.59) |
|  | Multiple | 1 (0.34) |
| **Material**<br>**availability**<br>**statement**<br>**(n=26)** | Personal or institutional | 5 (19.23) |
|  | Hosted by the journal | 18 (69.23) |
|  | Online third party | 1 (3.85) |
|  | Upon request | 2 (7.69) |
|  | Material were downloaded and accessible | 17 (65.38) |
|  | Material could not be downloaded and were not accessible | 9 (34.62) |
| **Data**<br>**availability**<br>**statement**<br>**(n=41)** | Personal or institutional | 4 (9.76) |
|  | Supplementary information hosted by the journal | 31 (75.61) |
|  | Online third party | 0 (0.0) |
|  | Upon request | 6 (14.63) |
|  | Data could be accessed and downloaded | 31 (75.61) |
|  | Data could not be accessed and downloaded | 10 (24.39) |
| **Documented data with all raw material (n=31)** | Data files clearly documented | 31 (100) |
|  | Data files not clearly documented | 0 (0.0) |
|  | Data files contain all raw data used for study conduction | 11 (35.48) |
|  | Data files do not contain all raw data used for study conduction | 20 (64.52) |
| **Pre-registration statement listed (n=10)** | Pre-registration could be accessed | 8 (80) |
|  | Pre-registration could not be accessed | 2 (20) |
|  | Pre-registered on clinicaltrials.gov | 6 (60) |
|  | Other ¶ | 4 (40) |
| **Documented within pre-registration (n=8)** | Hypothesis was included and documented within the pre-registration | 3 (37.5) |
|  | Methods was included and documented within the pre-registration | 8 (100) |
|  | Analysis plan was included and documented within the pre-registration | 0 (0.0) |
| **Protocol available (n=4)** | Hypothesis was included within the protocol | 0 (0.0) |
|  | Methods was included within the protocol | 3 (75.0) |
|  | Analysis plan was included within the protocol | 3 (75.0) |
| **Analysis script available (n=4)** | Personal or institutional | 0 (0.0) |
|  | Supplementary information hosted by the journal | 4 (100.0) |
|  | Online third party | 0 (0.0) |
|  | Upon request | 0 (0.0) |
| ¶ Includes: Australian New Zealand Clinical Trials Registry (n=1), Iranian Registry of Clinical Trials (n=1), ISRCTN Registry (n=1), National Research Register (n=1) |  |  |

### Evidence Synthesis

Of the 167 publications included in our analysis, the majority were not cited by either a meta-analysis or systematic review (140/167; 83.83%). Furthermore, no publications were replicated studies of previously published articles.

## Discussion

Results from our study indicate that the current state of nephrology research is not inclusive of transparent and reproducible research practices. Few studies in our sample provided access to study protocols, materials, analysis scripts, or study data. These results mirror a broad investigation of 441 publications in biomedical sciences, in which only 1 provided access to study protocols, 0 provided raw data, and only 4 were replication studies^8^. In this study, we highlight 2 important indicators of transparency and reproducibility that were exceptionally deficient in our sample. When discussing these indicators, we outline possible solutions for both funders and journals, when possible, and also describe current efforts underway to promote such practices.

First, data sharing is high yield for analytic reproducibility of a previous study. Few investigators made data publicly accessible, which hampers such efforts. From a funding perspective, various institutes within the NIH have established data repositories for institute-funded investigations. Some institutes have been more dedicated to these efforts than others. The National Institute of Allergy and Infectious Diseases, for example, supports 8 data repositories, including Immune Tolerance Network (ITN) TrialShare, which makes data and analysis code underlying ITN-published manuscripts publicly available with the goal of promoting transparency, reproducibility, and scientific collaboration^9^. The NIDDK funds a central repository that contains a biorepository that gathers, stores, and distributes biological samples from studies in addition to a data repository that stores data for all NIDDK grant research projects^10^. This central repository is important for the NIDDK, as their data sharing policy lists out various timelines for expected data to be deposited depending on the study design. The data sharing policy goes further to explain that all data will eventually become publicly available to increase its use and analysis in subsequent studies^11,12^. Furthermore, some private foundations require data sharing, such as the Bill and Melinda Gates Foundation^13^. Given that funders are able to impose such requirements, they have great influence on whether and how study data are made available. From a journal perspective, we selected 3 journals which had the highest h5-index rankings in Google Scholar’s urology and nephrology category (after excluding urology journals) to provide the basis for discussion. The *Journal of the American Society of Nephrology* (*JASN*) subscribes to the ICMJE Data Sharing policy for clinical trials^14^. All manuscripts of clinical trials must submit a data sharing plan according to ICMJE standards^15^. Data from systems-level analyses (e.g., such as genomics, metabolomics and proteomics) must be deposited in appropriate publicly accessible archiving sites. We could find no information on data sharing on *Kidney International’*s (*KI*) instructions for authors page^16^. The *American Journal of Kidney Diseases* (*AJKD*) requires all clinical trials to provide a data sharing statement. The instructions for authors notes that, “at this stage *AJKD* does not have a particular data sharing expectation”^17^. While a limited sample, differences within these journals showcase that variation exists by the very entities that arbitrate what and how nephrology research is published. We recommend that nephrology journals consider moving beyond indifference and adopt stricter policies. While dissenting views exist^18^, we believe that data sharing ultimately serves the best interests of the public, who provides tax dollars to fund research, and trial participants who subject themselves to potentially harmful interventions for the public good and want their data to count^19,20^.

Second, protocols were seldom provided by the study authors among publications in our sample. Detailed protocols are necessary for subsequent investigators to reproduce an original study or for readers to evaluate any deviations that occurred following protocol development. The Health and Human Services Department issued a “final rule” for clinical trials registration and results information submission. This rule specifies how data collected and analyzed during the trial should be reported to ClinicalTrials.gov. Specific to protocols, the rule requires “submission of the full version of the protocol and the statistical analysis plan (if a separate document) as part of clinical trial results information^21^. Thus, federal statutes require protocol sharing for some clinical trials. Building on our comparison of nephrology journals, we evaluated the guidance for submitting authors concerning protocol publication. We failed to locate any information on the *KI* or *JASN* authorship guidelines in regards to protocol submission except in the case of brief reports^14,16^. The *AJKD* requires that clinical trials include a study protocol with dated changes for the confidential review process, but leaves the protocol publication to the author’s discretion^17^. Thus, a comparison of current journal requirements for publishing research protocols suggests, at most, that the decision to publish a protocol may be left to the individual authors. When protocols are required at the time of submission, only peer reviewers and editors have access to them. When protocols remain unpublished, studies are unable to be reproduced or critically inspected. On many occasions, methods sections within published reports are too concise to truly understand whether the research methodology was robust or whether critical errors were made over the course of the study^22^. Protocols, which can easily be provided as supplementary documents on journal websites or deposited to platforms such as Open Science Framework (https://osf.io/), are necessary to fill in these gaps. Another platform, Protocols.io^23^, has been developed specifically for publication of research protocols. We also note that protocols are most often associated with clinical trials; however, we suggest that protocols are also necessary for other study types, such as observational studies. Protocols, for example, can carefully prespecify *a priori* which confounding factors are to be included in regression models. Absent publication of protocols, it is not possible to know whether model adjustments were made *post hoc*.

### Strengths and Limitations

Our study had many strengths including taking a random sample from a broad swath of all nephrology literature. Using this sampling methodology, we increase the likelihood that our results are generalizable to the nephrology research community as a whole. Our methodology, which included data extraction by 2 investigators in a blinded, duplicate fashion, is the gold standard methodology in systematic reviews and is endorsed by the Cochrane Collaboration. We also made our study protocol, materials, and data publicly available to foster greater transparency and reproducibility. Regarding limitations, our sample is restricted to journals which are indexed in PubMed and published in English. We also surveyed publications over a fixed time period. These considerations should be taken into account when interpreting and generalizing our study’s findings.

### Conclusion

In conclusion, our study found that reproducible and transparent research practices are infrequently employed by the nephrology research community. Greater efforts should be made by both funders and journals, two entities that have the greatest ability to influence change. In doing so, an open science culture may eventually become the norm rather than the exception.

## References

1. Veronese FV, Manfro RC, Roman FR, et al. Reproducibility of the Banff classification in subclinical kidney transplant rejection. Clinical Transplantation. 2005;19(4):518–521. doi:10.1111/j.1399-0012.2005.00377.x

2. Piskunowicz M, Hofmann L, Zuercher E, et al. A new technique with high reproducibility to estimate renal oxygenation using BOLD-MRI in chronic kidney disease. Magn Reson Imaging. 2015;33(3):253–261.

3. Affret A, Wagner S, El Fatouhi D, et al. Validity and reproducibility of a short food frequency questionnaire among patients with chronic kidney disease. BMC Nephrol. 2017;18(1):297.

4. dkNET | Introduction. https://dknet.org/about/product_info. Accessed August 16, 2019.

5. dkNET | NIH Policy Rigor Reproducibility. https://dknet.org/about/NIH-Policy-Rigor-Reproducibility. Accessed August 16, 2019.

6. Enhancing Reproducibility through Rigor and Transparency | grants.nih.gov. https://grants.nih.gov/policy/reproducibility/index.htm. Accessed August 16, 2019.

7. RFA-GM-19-001: Methods to Improve Reproducibility of Human iPSC Derivation, Growth and Differentiation (SBIR) (R44 Clinical Trial Not Allowed). https://grants.nih.gov/grants/guide/rfa-files/RFA-GM-19-001.html. Accessed August 16, 2019.

8. Iqbal SA, Wallach JD, Khoury MJ, Schully SD, Ioannidis JPA. Reproducible Research Practices and Transparency across the Biomedical Literature. PLoS Biol. 2016;14(1):e1002333.

9. Sign In: /home. https://www.itntrialshare.org/login/home/login.view?returnUrl=%2Fproject%2Fhome%2Fstart.view%3F. Accessed August 16, 2019.

10. Hornik K, Others. The r FAQ. 2002. https://repository.niddk.nih.gov/faq/.

11. Rasooly RS, Akolkar B, Spain LM, Guill MH, Del Vecchio CT, Carroll LE. The National Institute of Diabetes and Digestive and Kidney Diseases Central Repositories: a valuable resource for nephrology research. Clin J Am Soc Nephrol. 2015;10(4):710–715.

12. Policies for Clinical Researchers | NIDDK. National Institute of Diabetes and Digestive and Kidney Diseases. https://www.niddk.nih.gov/research-funding/human-subjects-research/policies-clinical-researchers. Accessed August 16, 2019.

13. Open Access Policy. Bill & Melinda Gates Foundation. https://www.gatesfoundation.org/How-We-Work/General-Information/Open-Access-Policy/Page-2. Accessed August 12, 2019.

14. Author Resources | American Society of Nephrology. https://jasn.asnjournals.org/content/authors/ifora. Accessed August 16, 2019.

15. Taichman DB, Sahni P, Pinborg A, et al. Data Sharing Statements for Clinical Trials: A Requirement of the International Committee of Medical Journal Editors. Ethiop J Health Sci. 2017;27(4):315–318.

16. Elsevier. Guide for authors - Kidney International - ISSN 0085-2538. https://www.elsevier.com/journals/kidney-international/0085-2538/guide-for-authors. Accessed 1 August 16, 2019.

17. AJKD Information for Authors & Journal Policies. https://sites.google.com/site/ajkdinfoforauthors/. Accessed August 16, 2019.

18. Longo DL, Drazen JM. Data Sharing. N Engl J Med. 2016;374(3):276–277.

19. Alliance - Resolution to Share Legacy Clinical Trial Data. https://www.allianceforclinicaltrialsinoncology.org/main/public/standard.xhtml?path=%2FPublic%2FPAC-Resolution. Accessed August 16, 2019.

20. Aitken M, de St Jorre J, Pagliari C, Jepson R, Cunningham-Burley S. Public responses to the sharing and linkage of health data for research purposes: a systematic review and thematic synthesis of qualitative studies. BMC Med Ethics. 2016;17(1):73.

21. Health and Human Services Department. Clinical Trials Registration and Results Information Submission. Federal Register. 2016;81:64981–65157. https://www.federalregister.gov/d/2016-22129.

22. Steward O, Popovich PG, Dietrich WD, Kleitman N. Replication and reproducibility in spinal cord injury research. Exp Neurol. 2012;233(2):597–605.

23. About protocols.io. https://www.protocols.io/about. Accessed August 16, 2019.

