## Supplemental Table 1 for "An evaluation of Nephrology Literature for Transparency and Reproducibility Indicators: Cross-sectional Review"

**Supplemental Table 1:** Sample characteristics of analyzed sports medicine publications

| ***Characteristic*** | | ***Variables*** |
| --- | --- | --- |
|  | | **No. (%)** |
| **Study subjects (n=295)** | Animals | 39 (13.22) |
|  | Humans | 145 (49.15) |
|  | Both | 2 (0.68) |
|  | Neither | 109 (36.95) |
| **Country of journal publication (n=295)** | US | 182 (61.69) |
|  | UK | 43 (14.58) |
|  | Switzerland | 22 (7.46) |
|  | Netherlands | 13 (4.41) |
|  | India | 9 (3.05) |
|  | Germany | 8 (2.71) |
|  | Brazil | 7 (2.37) |
|  | Canada | 5 (1.69) |
|  | Italy | 2 (0.68) |
|  | Japan | 2 (0.68) |
|  | Iran | 2 (0.68) |
| **Country of corresponding author (n=295)** | US | 97 (32.88) |
|  | China | 26 (8.81) |
|  | Japan | 21 (7.12) |
|  | UK | 12 (4.07) |
|  | Australia | 11 (3.73) |
|  | Germany | 11 (3.73) |
|  | Canada | 10 (3.39) |
|  | Turkey | 10 (3.39) |
|  | France | 9 (3.05) |
|  | Brazil | 8 (2.71) |
|  | Netherlands | 8 (2.71) |
|  | Italy | 7 (2.37) |
|  | South Korea | 6 (2.03) |
|  | Unclear | 6 (2.03) |
|  | India | 5 (1.69) |
|  | Spain | 4 (1.36) |
|  | Other † | 44 (14.92) |
| **Most recent impact factor year (n=295)** | 2018 | 276 |
|  | Not found | 19 |
| **5 Year impact factor (n=276)** | Median | 3.232 |
|  | 1st quartile | 2.053 |
|  | 3rd quartile | 7.065 |
|  | Interquartile range | [2.053-7.065] |
| **Most recent impact factor (n=276)** | Median | 3.013 |
|  | 1st quartile | 1.971 |
|  | 3rd quartile | 6.653 |
|  | Interquartile range | [1.971-6.653] |
| Abbreviations: CI, Confidence Interval.  † Includes: Austria n=1, Belgium n=4, Chile n=1, Croatia n=1, Cyprus n=1, Czech Republic n=1, Denmark n=1, Egypt n=2, Finland n=1, Greece n=1, Iran n=4, Ireland n=1, Israel n=1, Lebanon n=1, Lithuania n=1, Malaysia n=2, New Zealand n=1, Nigeria, n=1, Norway n=2, Poland n=1, Portugal n=1, Saudi Arabia n=3, Scotland n=1, Sweden n=3, Switzerland n=3, Taiwan n=3, Thailand n=1 | | |
